# Probing The Edge Of Synchronization: Cortical Activity Transiently Slips Into Slow Waves In Behaving Monkeys

**DOI:** 10.1101/2024.09.20.613889

**Authors:** Antonio Pazienti, Andrea Galluzzi, Pierpaolo Pani, Stefano Ferraina, Maurizio Mattia

**Affiliations:** Natl. Center for Radioprotection and Computational Physics, Istituto Superiore di Sanità, 00161 Rome, Italy; Dept. of Physiology and Pharmacology, Sapienza University, 00185 Rome, Italy

**Keywords:** Slow waves, Cortical networks, Critical dynamics, Sleep-wake cycle, Delayed-reaching task, Spiking neuron models

## Abstract

Brain networks oscillate between sleep and wakefulness, following circadian rhythms. Theoretical models suggest distinct phases within this cycle, separated by a critical point where long-range activity patterns emerge, an advantageous condition for information processing in cortical networks. However, the exact nature of this critical dynamics remains elusive. A key question is whether the brain operates at this critical point during cognitive tasks or only during resting wakefulness. Here, we analyzed neural signals from the premotor cortex (PMC) of two macaque monkeys engaged in a delayed-reaching task and under drug-induced unconsciousness. We found evidence of criticality during resting periods at the end of behavioral trials in the awake state. This scale-free activity appeared as coordinated traveling waves, like those observed during anesthesia. As predicted by spiking networks models, activity-dependent adaptation influences wave size, supporting the hypothesis that the PMC operates near a synchronization phase transition while avoiding it during active behaviors.

## Introduction

The mammalian brain is a complex system perfectly designed to be capable to collect information from the environment as sensory stimuli, organize this information for forming internal representations, and coordinate the muscular output to fulfil goals (Harris and Thiele, 2011). The healthy brain can be found in various global states such as active wakefulness, rest, and sleep (Gervasoni et al., 2004); the dynamics characteristic of these states allows the brain to express its entire performance, preserve energy, and at the same time consolidate and replenish resources (Buzsáki, 2004; Diekelmann and Born, 2010).

In the last decades, the brain has been associated to a self-organized critical system, whose peculiar property is to intermediately store incoming energy and to intermittently release it (Christensen and Moloney, 2005). Generally, the response of a system at the critical point is uncorrelated to the perturbation it receives and can span an arbitrary number of temporal and spatial scales, limited only by its own size. Critical activity has been mostly associated with the awake state, where computational power is maximally needed (Bellay et al., 2015; Ma et al., 2019; Meshulam et al., 2019). Assessing criticality in the brain is tricky though, as it requires highly spatially sampled signals; indeed, subsampling was shown to easily lead to erroneous assessments of criticality (Priesemann et al., 2009; Priesemann, 2014; Levina and Priesemann, 2017; Wilting and Priesemann, 2019).

Recently, evidence has accumulated that the brain activity resembles less that of a classical sandpile system, rather crossing the critical boundary to synchronization, separating an active desynchronized —that is, with a high, irregular neuronal activity, typical of attentive wakefulness—from an isolated synchronized state—that is, with an activity switching between silence and strong coordinated activations, typical of sleep – namely the slow-wave activity (SWA) (Di Santo et al., 2018; Fontenele et al., 2019). In this view, the relevant state transition is between the conscious and the unconscious state, and approaching the critical point is directly related to a change in the global state of the brain (Sanchez-Vives et al., 2017; Levenstein et al., 2019; Tort-Colet et al., 2021). This hypothesis does not contradict the data showing that given the global state of the brain, its subnetworks can be dwelling in different local states (Huber et al., 2004; Mahowald and Schenck, 2005; Nir et al., 2011; Vyazovskiy et al., 2011). However, a detailed description of the dynamics of the relationship between the occurrence of criticality and the behavioral activity remains to be elucidated.

With the present work we question these views and address what kind of criticality the brain—or portions of it—is nearing during a sensorimotor task performance. Ultimately, we come to measurable quantities predictive of vicinity to the critical point. We used millisecond-precise and highly spatially-resolved neural signals from a relevant portion of the macaque brain both while performing a delayed-reaching task and during sleep-like, anesthetics-induced unconsciousness. We found that the awake brain is *repetitively* nearing a synchronization phase transition separating the active and the synchronized state. This dynamics emerged during short resting periods at the end of each trial and was characterized by the occurrence of neuronal avalanches and of coordinated neuronal activations traveling in space—assimilable to sleep-like slow waves. However, during the active task performance, the dynamics resulted far from the critical point. Importantly, we also designed a spiking network model, whose dynamics was guided to cross the critical point separating the desynchronized and synchronized global states. The prediction of the model that activity-dependent adaptation determines the size of avalanches (and slow waves) was confirmed in the experimental data, thereby strengthening the conclusion that synchronous activations in rest and slow waves under anesthesia were expression of the same phenomenon.

Our results unveil the details of a basic albeit profound mechanism with which the brain transitions between different global states, thus shedding new light on the comprehension of the brain’s fundamental homeostatic functions.

## Results

### Premotor cortex escapes equilibrium during a motor task

We performed multi-electrode intracortical recordings (96-channels Utah array, Blackrock Neurotech, USA) from the dorsal premotor cortex (PMd) of two male rhesus monkeys (*Macaca mulatta*) (Figure 1A; see Methods) while both monkeys performed a delayed-reaching task (Weinrich and Wise, 1982; Crammond and Kalaska, 2000). To receive a reward, animals were trained to move their arm towards a target presented on a touchscreen following a Go signal after a variable-duration delay. From the electrophysiological recordings we extracted both the local field potential (LFP) and the spectral-derived multi-unit spiking activity (also referred to as normalized firing rate (FR), see Methods) (Supèr and Roelfsema, 2005; Stark and Abeles, 2007; Mattia et al., 2010) of assemblies of neurons.

**Figure 1.**
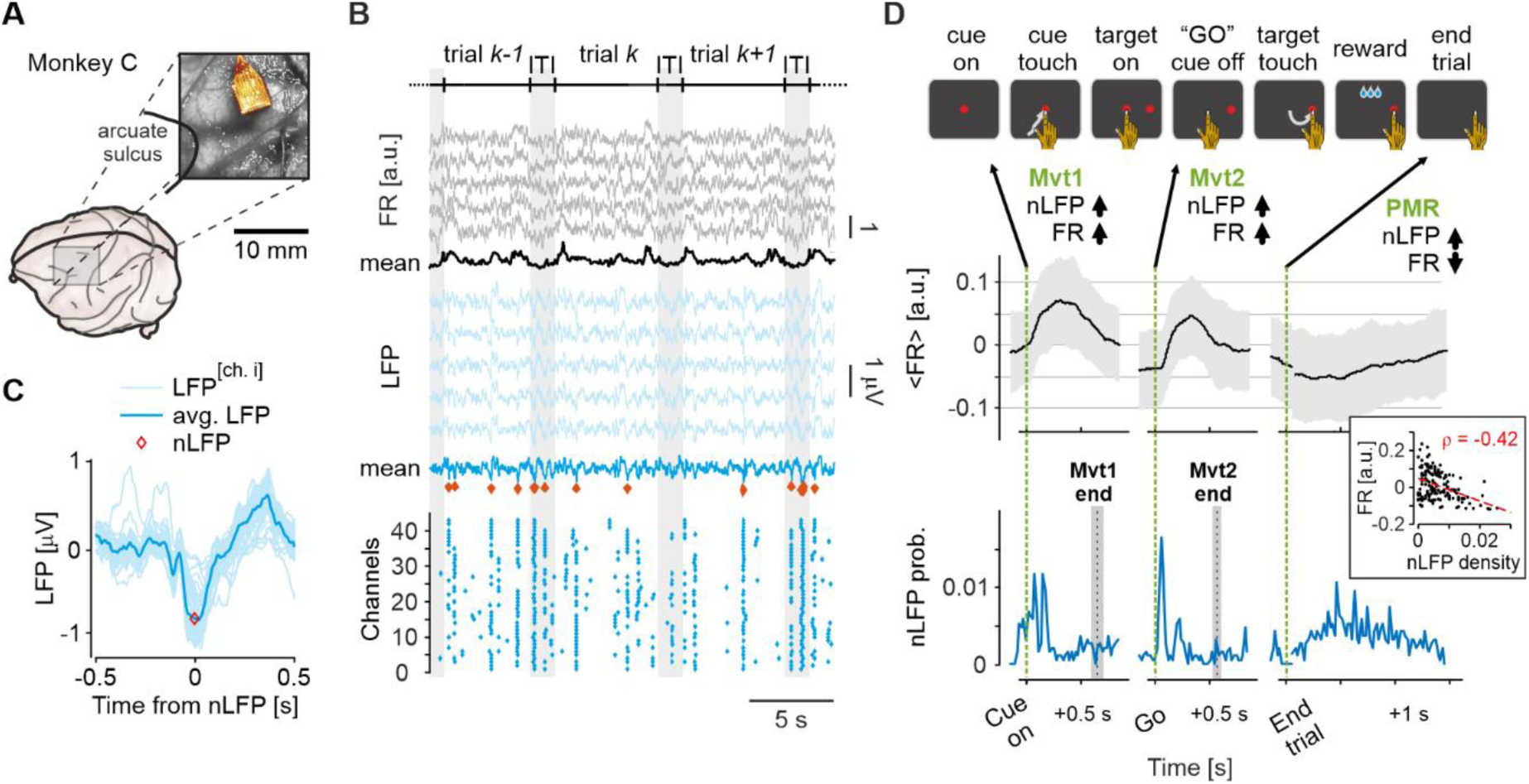
Large LFP fluctuations occurring after movement execution take PMd out of equilibrium. **A)** Position of the multi-electrode array (96-channel Utah array) on the brain surface (monkey C). **B)** In a subsample of recording channels, normalized firing rate (FR) (top), local field potential (LFP) (middle), and raster plot of the nLFPs channel-wise (bottom). Red diamonds: nLFPs detected from the average LFP signal. Vertical grey shaded periods: inter-trial intervals (ITI). Average ITI duration: 1.50 ± 0.02. **C)** Representative cut-out of the LFP around the nLFPs for all channels and their average (darker line). **D)** Identification of the interval of interest for the analysis of criticality during periods of reduced firing activity. The interelectrode average FR (± SD) (middle) and the probability density (bottom) of the nLFPs (as in panel B) are plotted in intervals of different duration aligned to key task events (shown on top). For this monkey (C), the ITI – here indicated as PMR (post movement resting period) shows the lowest average FR. In the same period, nLFPs occur with the highest frequency. Mvt1 and Mvt2 indicate, respectively, the two epochs where the monkey generate a movement. Mvt1: first movement in the trial, after the Cue onset the arm is moved to touch the cue. Mvt2: second movement in the trial, after the “Go” signal the arm is moved to touch the peripheral target. Inset: anti-correlation between FR and nLFP density across all the ITIs (*ϱ* = – 0.42, *P* < 10^−7^) of the recording session.

We collected these signals in order to investigate the synaptic input and the output of our system and relate its dynamics in the state space to the behavior. Indeed, the scaling properties of fluctuations of the LFP as a function of their size have been used to associate the brain activity to a critical state (Petermann et al., 2009). As a proxy to these fluctuations, we extracted the negative peaks of local field potentials (nLFP) both from all PMd recording channels and from the averaged signal (Fig. 1B, C). Negative peaks of the mean LFP co-occurred with subsets of single-channel nLFPs of variable sizes (Figure 1B, bottom). As expected from PMd, the movement generation was systematically accompanied by large increases of the FR (Figure 1D, middle) and sudden changes in the density of nLFPs (Figure 1D, bottom). The latter LFP fluctuations were not movement-related (see end-of-movement labels on Fig. 1D), but clearly time-locked to external, task-related cues—e.g., target appearance—compatibly with the hypothesis that they can be associated to the precise timing of the synaptic input from other brain areas (Buzsáki et al., 2012). Interestingly however, we also observed a persistent and significantly high presence of nLFPs in the resting periods, just after the execution of the last movement in the trial, possibly including part of the inter-trial intervals (Figure 1D, bottom). In these periods, both arm movements and sensory stimuli were absent, thus putting the local network in a condition of relative isolation and rest. Importantly, in these instances high fluctuations co-occurred with a very low spiking activity of individual assemblies, the two quantities being significantly anti-correlated (Fig. 1D, inset). As FR reductions have been linked with evidence of critical dynamics (Ma et al., 2019), we focused our subsequent analysis on these periods of low FR, which we will refer to as post-movement resting (PMR), as a candidate period to get closer to a brain state transition during wakefulness.

### Footprints of criticality during periods of reduced firing activity

We have identified periods of high synaptic and low spiking activity unrelated to any incoming visual stimuli or movement generation. These instances may underlie the temporary permanence of the network collective dynamics in the vicinity of a critical point (Ma et al., 2019). To address this possibility, we thoroughly examined the temporal relationship between the LFP fluctuations and the spiking activity of the explored assemblies of neurons.

In the PMR epochs, we observed a sudden and significant increase of FR preceding the nLFP occurrence (Fig. 2A) by 90-140 ms (monkey P: 141 ms, monkey C: 91 ms; see Methods). This sharp increase in spiking activity could be unveiled only by aligning on the nLFPs, as it is not locked to the external trial cues (cf. Fig. 1). However, the jump in spiking activity involves most of the channels (Figure 2B; yellow circles). Importantly, the intensity of the FR increase correlated with the amplitude of the nLFPs troughs (not shown), and such an increase was not present when taking random times in the same PMR epochs (Fig. 2A, gray lines). We will refer to this coordinated increase in FR, as ‘sharp upward transitions’ (SUT) (see also (Mattia et al., 2013)).

**Figure 2.**
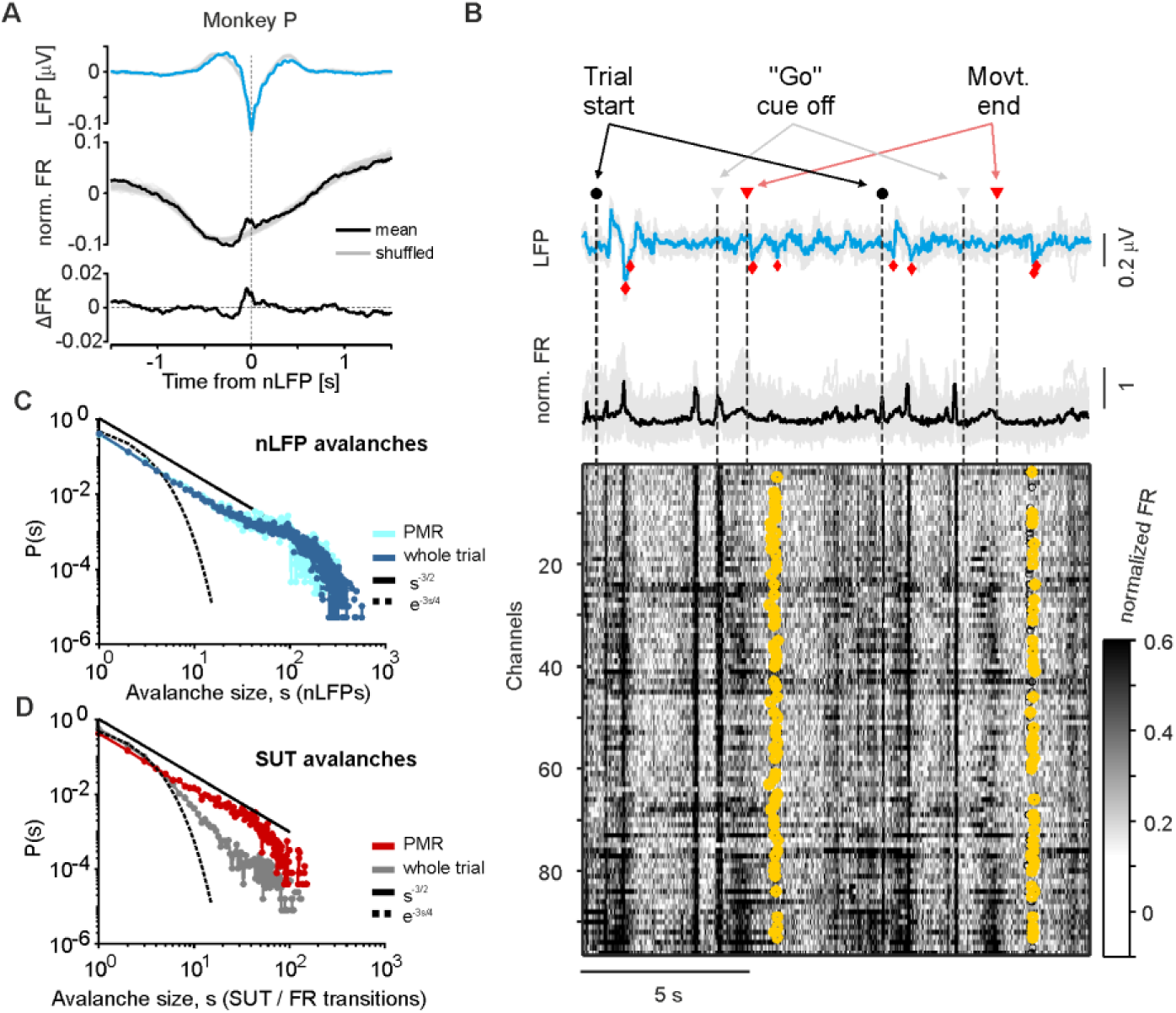
Both nLFPs and SUT show signs of criticality during rest. **A)** Average LFP (top, blue) and average normalized FR (middle, black) across the electrode array in the PMR epoch for monkey P. Grey traces: average LFP around the same number of nLFP but randomly chosen in the whole trial (top) and mean FR across 50 random samplings of the same time interval (middle). Bottom: difference between the mean and the shuffled FR. **B)** Representative average LFP (blue, top) and average FR (black, middle) in two consecutive trials. Grey traces are the corresponding single-channel LFP and FR. Bottom: raster activity of the FR for all the channels. Yellow circles: SUT detected at single-channel level in the PMR. **C-D)** Distribution of avalanche size singled out from nLFPs (C) and SUT (D) grouped by PMR and whole-trial epochs. Black full and dashed line show the -3/2 power-law and the gaussian regimes, respectively.

We then investigated the possible scale-free organization of the neuronal dynamics by computing neuronal avalanches both in PMR epochs and in the remaining portions of the task. We first computed the widely used distribution of nLFP size across channels (Petermann et al., 2009). The distribution of avalanche size for nLFPs in our data is shown in Fig. 2C. The distribution resulted to be roughly scale-free both in the PMR epoch and—importantly—in the rest of the task (Fig. 2C). Subsequently, we estimated the scale-free features using the single-channel SUTs. We found a scale-free distribution of SUTs only during PMR, whereas when considering the entire trial, the distribution was clearly subcritical (Fig. 2D).

These findings reveal that during periods of rest and low spiking activity, while the animal was actively engaged in the task, the cortical network exhibits collective neuronal events—involving both the synaptic input and the output of the neurons—that are scale invariant over multiple orders of magnitude. In the following, we will characterize this possible signature of the system approaching a critical state and detail, spatially and temporally, what type of network activity is involved.

### SUTs underlie travelling waves in PMd during resting

The collective network activations we have documented turned out to lack any characteristic scale, to occur during periods of low spiking activity and to follow synaptic fluctuations. This phenomenon could be the signature of the system approaching a critical point. We therefore asked whether the network activity displayed any spatiotemporal pattern usually lacking during. In other words: do the activations associated with the SUT entrained the neurons simultaneously, in a random order or with a certain spatial and/or temporal specific characteristic? For this purpose, we analyzed the exact time of occurrence of the SUTs for each channel and their spatiotemporal organization on the cortical surface.

In order to characterize the participation of the individual channels to the SUT in the PMR, we extracted the time of activation of every channel (Fig. 3A; the relative value composes the time-lag matrix, see Methods and (Mattia et al., 2010; Capone et al., 2019; Pazienti et al., 2022)) and grouped similar patterns of activation referring to them as ‘modes’ (Dasilva et al., 2021; Pazienti et al., 2022). Surprisingly, the channels showed a wave-like coordination in time and space, forming spatiotemporal patterns reminiscent of slow-wave cortical activity. The wavefront propagated across the grid at a velocity of about 15 mm/s [17.9 ± 0.3 and 12.7 ± 0.2 mm/s, for monkeys C and P, respectively] (Fig. 3B, left). The spatiotemporal patterns we obtained showed a clear structure, with closely located channels activating first, and the rest of the channels progressively being recruited (Fig. 3B).

**Figure 3.**
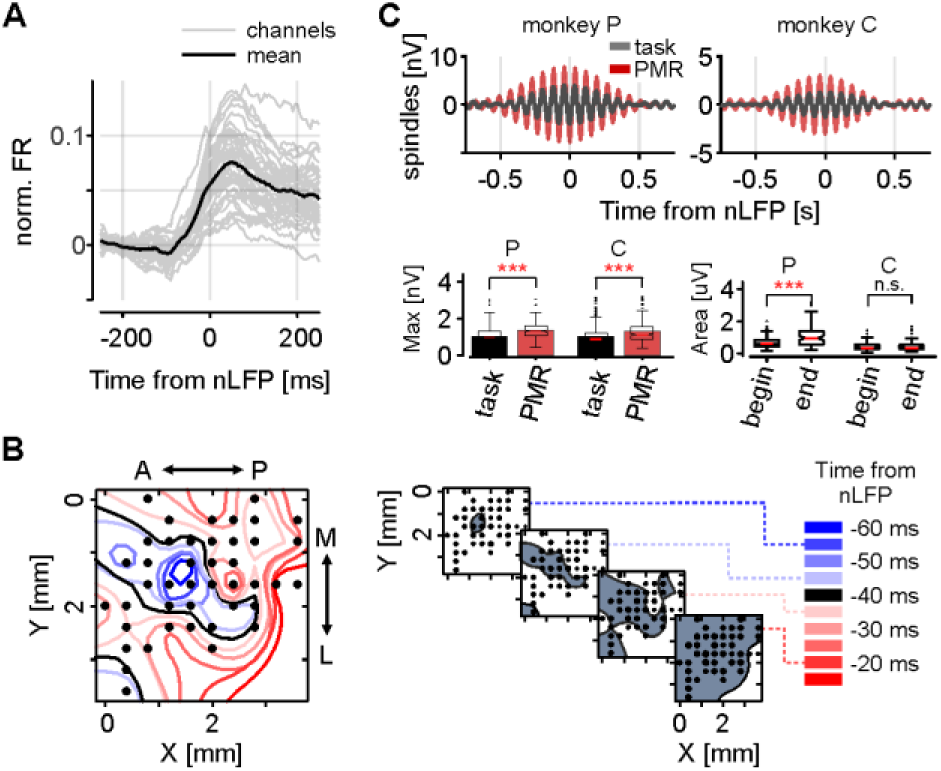
Signature of slow-wave dynamics in PMd during the task execution. **A)** SUT averaged across all detected transitions (grey) and average of all channels (black). **B)** Left, time of activation of each channel on the electrode grid (recording channels are shown as black dots in their actual physical location) relative to the nLFP. Shades of blue show channels anticipating the SUT, whereas shades of red indicate channels following the transition. (Right) Snapshots of the propagating increase of FR on the cortical surface for four different time delay with respect to the nLFP occurrence. As a reference, times from the average nLFP are shown. **C)** nLFP-centered spindle (12-14 Hz)-component of the LFP for monkeys P and C in the PMR epochs (red) and in the rest of the trial (grey). Bottom: pooled data of the spindle maxima during the same periods (left) and spindle area during the task in the first vs. the last 100 trials (right) for both monkeys.

A neurophysiological signature of brain activity during sleep is the presence of so-called spindles, having a strong component at frequencies around 12 Hz (Loomis et al., 1935; Fernandez & Lüthi, 2020). If our system was approaching the wake-sleep transition in the PMR epochs, we would expect the fluctuations of the electrophysiological signals to become stronger. We therefore looked for these signatures during the experimental session. The nLFP amplitude at the end of the session (last 100 nLFPs) was significantly higher than the first 100 nLFPs for both monkeys (not shown, *p* = 8.6 × 10^−8^ for monkey P and *p* = 3.3 × 10^−5^ for monkey C). Moreover, the size of the spindles was significantly higher in the last vs. the first 100 trials (Fig. 3C, bottom right; significant in one monkey and as a trend in the other monkey; *p* = 5.9 × 10^−6^ for monkey P and *p* = 0.26 for monkey C). Surprisingly, we found a similar strong increase of spindle amplitude when comparing the PMR with respect to the rest of the trial during the whole session (Fig. 3C, bottom left), starting 3-400 ms before the nLFP events, and lasting more for about 300-500 ms (Fig. 3C).

Together, these results show that during short instances of resting following a task execution the network is subject to the emergence of spatiotemporally coordinated patterns spreading over several millimeters resembling the activity typically observed during sleep, and this observation being also supported by other neurophysiological signatures.

### Slow waves in PMd activate neuronal assemblies similarly to SUT in resting

We have documented the activation of large groups of neurons in a coordinated manner—the phenomenon we termed the SUTs—during resting periods of the task. This reminded us of Down-to-Up transitions during sleep- or anesthesia-induced slow oscillations (SO) (Sanchez-Vives and McCormick, 2000; Sanchez-Vives et al., 2017; Sanchez-Vives, 2020). Indeed, the precisely locked co-occurrence of nLFPs, Down-to-Up transitions and amplitude modulations of the spindle activity represents a clear signature of slow-wave activity (SWA), as documented both during natural sleep and under anesthesia (Vyazovskiy et al., 2009; Fernandez and Lüthi, 2020). To investigate parallels between these phenomena we induced stable anesthetized states in the same animals by injecting ketamine i.m. (see Materials and Methods for details). Reliable slow oscillations quickly arose in both monkeys, with the FR switching between periods of almost quiescence (Down states) and periods of relatively high-firing activity (Up states) (Fig. 4A). We then extracted the nLFPs and the times of increase in FR (Down-to-Up transitions) as in the previous sections. The Down-to-Up transitions of the FR were consistently phase-locked to nLFPs (Fig. 4A, middle). This is particularly apparent when inspecting the nLFPs-centered average FR. A rather stereotyped collective Down-to-Up transition is clearly visible, which anticipates the LFP trough (top and middle panels in Fig. 4B). The nLFP-centered activity also showed a clear modulation in the spindle frequency band (12-14 Hz, Fig. 4B, bottom), in a striking similarity with the result during the PMR epochs (cf. Fig. 3C).

**Figure 4.**
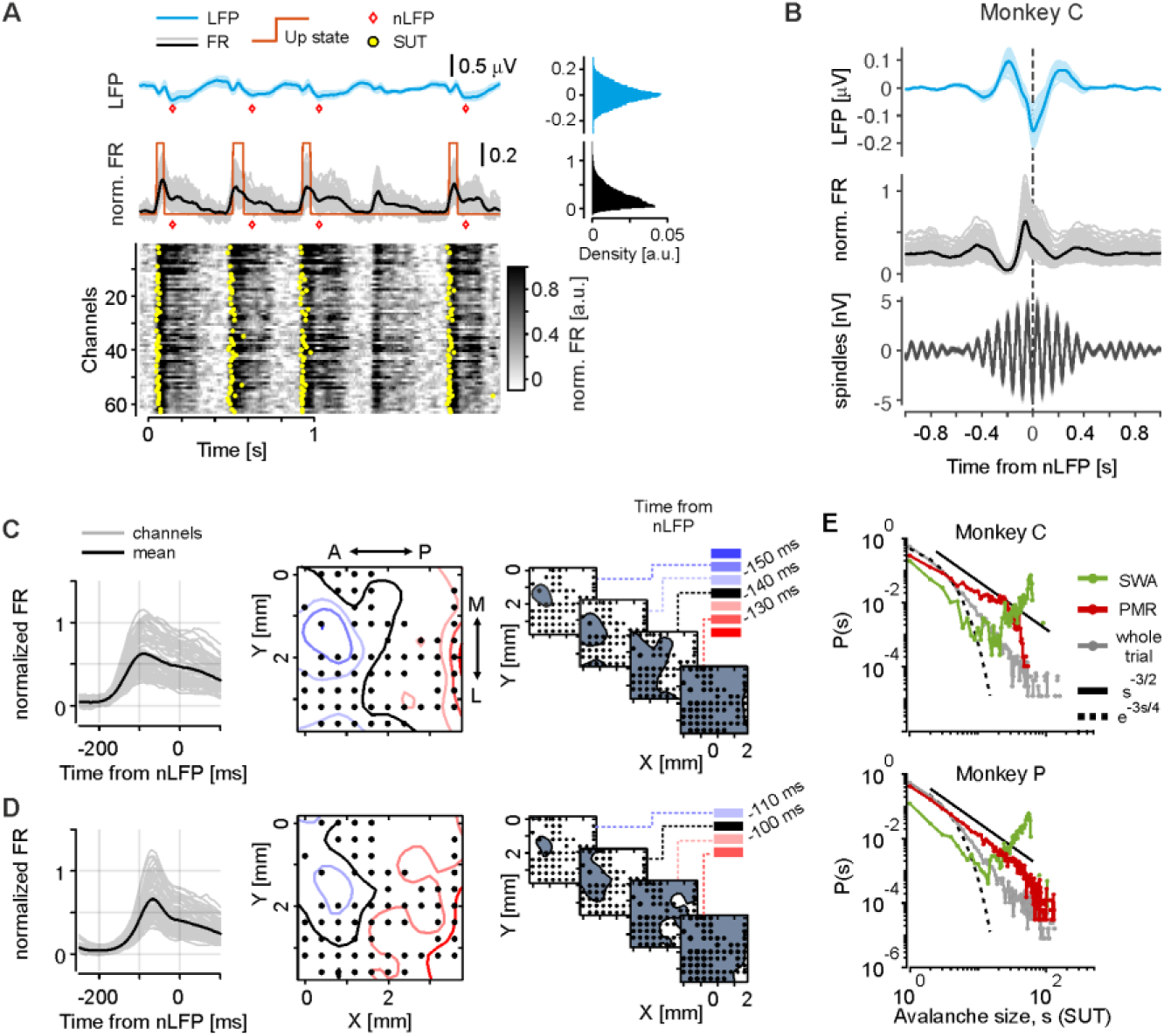
Slow-wave activity and travelling waves in PMd during unconscious state. **A)** Representative LFP (top), average (middle) and single-channel FR (bottom) during ketamine anesthesia for monkey C. Red diamonds indicate occurrence of nLFPs calculated from the average LFP. Red lines show detected Down-to-Up transitions; yellow dots indicate SUT occurrences. Top right: distribution in time of the waveforms on the left. **B)** Average nLFP across channels (top), nLFP-centered average FR (middle) and nLFP-centered spindles during anesthesia. **C)** From left to right, average FR transitions for each channel (grey) belonging to a group of slow waves, average of all channels (black). Negative times indicates anticipation with respect to other channels, the zero-value (black curve) being arbitrarily set to have a clear visualization; snapshots of the activity level for four different time delay with respect to the nLFP occurrence. **D)** Same as C), for the transitions belonging to another mode of slow waves. **E)** Probability of neuronal avalanches for SUT of the FR during SWA (green), PMR (red), and the whole trial (grey) for monkey C (top) and P (bottom). Black full and dashed line show the -3/2 power-law and the Gaussian regimes, respectively.

The spiking activity propagated (Fig. 4C) from the leading channels to the rest of the cortical area at a mean velocity of 11.6 ± 0.1 and 14.6 ± 0.6 mm/s for monkey P and C, respectively (*n* = 380 and *n* = 276 detected waves for monkey P and C, respectively). Next, we looked for neuronal avalanches, using the same procedure as with the SUT in the awake state. The size distribution shows a marked excess of large avalanches (the so-called king avalanches) for both monkeys (green curves in Fig. 4E). This can be explained by the fact that SWA avalanches mostly involved the entirety of the cortical field.

These results show that under ketamine anesthesia area PMd features sleep-like slow oscillations, in which organized spatiotemporal patterns travel across several millimeters. As observed in the resting epoch of the task (PMR epoch), the Down-to-Up transitions consistently preceded the occurrence of nLFPs and were accompanied by modulation of activity in the spindle frequency band. These results strengthen the similarity between the network patterns we found during resting and the unconscious state.

### Network model of spiking neurons at the edge of synchronization

The spatiotemporal patterns detected in the PMR suggested that during the execution of a delayed-reaching task neuronal the PMd network transitorily approaches the border separating the active and the oscillatory, sleep-like state. To further investigate this hypothesis, we simulated a network of leaky integrate-and-fire (LIF) model neurons connected with grid-like topology mimicking the recording arrays used in the experiments. The network was composed of 13×13 modules, each with 1,000 excitatory and 250 inhibitory LIF neurons, connected with probability decaying with the distance on the grid (Fig. 5A, and Materials and Methods). Within each module, the connection probability was 80%, the excitatory neuronal pool received an external current *I_ext_* and was subject to adaptation regulated by the parameter *g_a_* (Mattia et al., 2013). This kind of neuronal networks is a well-established model that can be set to produce various regimes of spiking activity (sketched in Fig. 5A, bottom), whose phase space can be explored by operating on the order parameters *I_ext_* and *g_a_*, namely the excitation-adaptation (E-A) bifurcation plane (Mattia & Sanchez-Vives, 2012).

**Figure 5.**
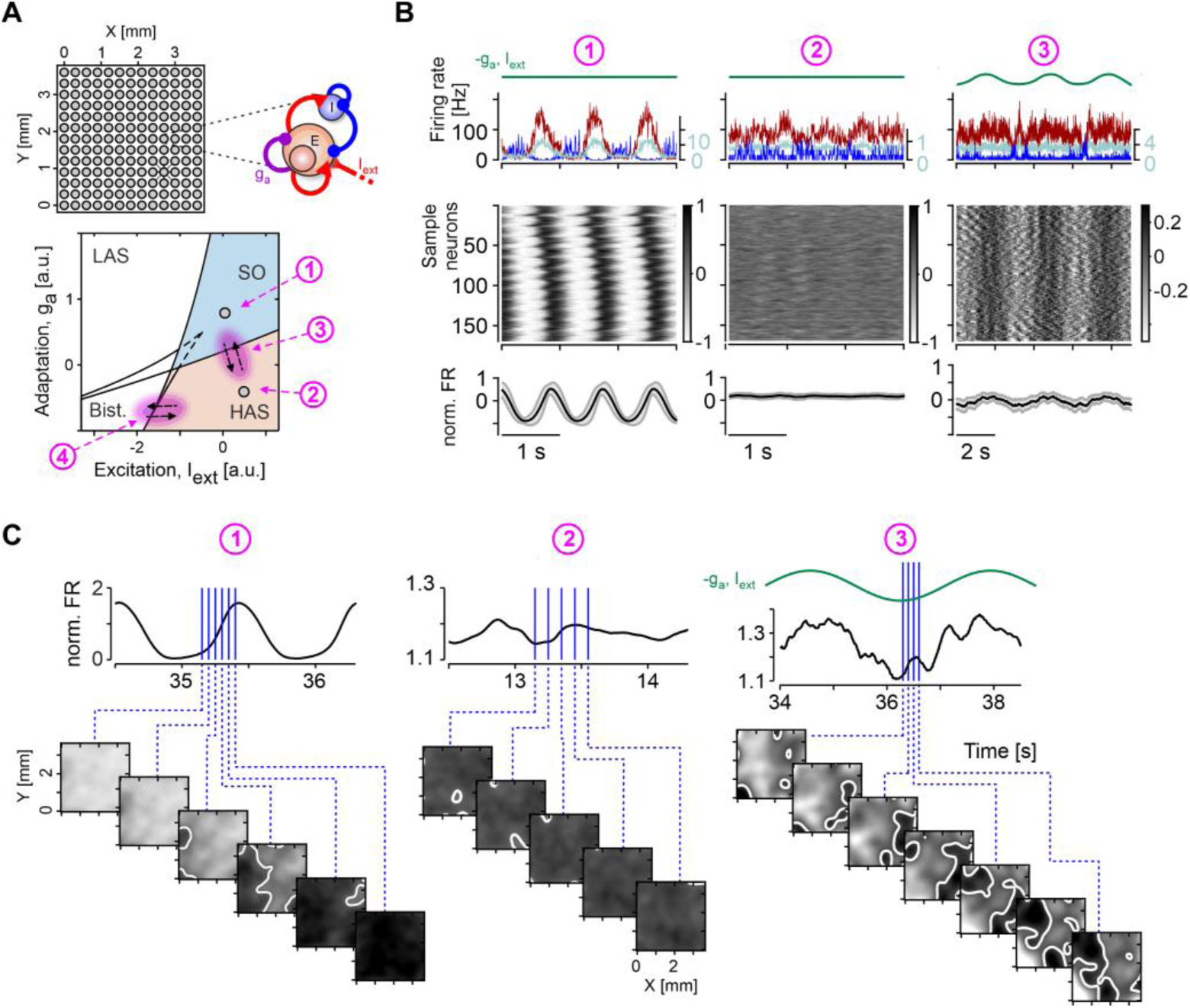
Network model dynamically exploring the edge of synchronization. **A)** Top, sketch illustrating the *in silico* network structure. The network is a two-dimensional grid composed of 13 × 13 modules, that communicate only through projecting excitation (see Materials and Methods). Internally, each module consists of 1,000 excitatory and 250 inhibitory LIF neurons connected with probability equal to 0.8. Excitatory neurons receive an additional external current (*I_ext_*) and are subject to spike-frequency adaptation (*g_a_*). Bottom, qualitative bifurcation diagram for a single module (Mattia & Sanchez-Vives, 2012). By changing the excitation *I_ext_* and the adaptation *g_a_* the dynamical regime of the network ranges from single to bistable point attractor (LAS, HAS and Bist. regions) including also stable limit cycles (SO region). Pink area and numbers indicate the phases explored in this work and detailed in panels B and C and in Fig. 6. **B)** Activity of neuronal populations at different network states. Top, value of *I_ext_* for simulations 1-3 and average firing rate of neuronal populations. Blue: inhibitory neurons; pink: excitatory background neurons (right y-axis); red: excitatory foreground neurons. Middle and bottom, normalized and average FR of the excitatory foreground population (see Methods). Example simulations are shown for the SO regime (left), in the high-firing asynchronous state (HAS, center) and back and forth across the critical line between the former regimes. For the latter simulation, *I_ext_* is sinusoidally modulated with a cycle period of 3 seconds (Top green traces) to model trial alternation in the monkey awake state. **C)** Top, normalized FR in a smaller time segment than in (B). Bottom, snapshots of the firing rate on the grid taken at several time points. In the SO state (left), the snapshots are taken during the Down-to-Up transition. In the simulation across the critical line (right), the snapshots are taken around the point closest to the SO region, where the emergence of spatiotemporal patterns across the grid is observed. LAS, low-firing asynchronous state.

Four dynamical phases of interest can be expressed by the excitation-adaptation parameters: 1) slow oscillations (SO, corresponding to the anesthetic-induced SWA in the experimental data); 2) irregular, high-firing asynchronous (HAS); 3) back and forth drifting across the critical line separating the two previous states and 4) a similar back and forth across the critical line separating the bistable and the HAS states (Fig. 5A, bottom). The network was capable to qualitatively reproduce the asynchronous and the oscillating spiking activity characteristic of the awake and the anesthetized state, respectively (Fig. 5B, left and middle). Importantly, in the SO state the network generated spatiotemporal patterns travelling across the grid that closely resembled the patterns we observed during anesthesia (cf. Fig. 5C, left with Fig. 4D, E).

In the condition 3—repeatedly crossing the critical point separating HAS and SO regions—the dynamics of the network naturally switched between asynchronous and synchronous spiking activity. Following each “cycle”, in which the FR decreased (cf. time values shortly after time=36 s in Fig. 5C, top right) the vicinity to the SO regime lead to brief collective activations that marked the initiation of the oscillatory activity (however immediately aborted as the parameters’ change brought the network back the the asynchronous state). This effect resulted in both sharp increases of the FR and coordinated spatiotemporal patterns between the nodes of our network. Critically, the patterns closely resembled the ones we observed experimentally during the PMR epochs of the task (cf. Fig. 5C, right with Figs. 2A and 3D).

In this regime we expected that, when the system crossed the critical line, the intrinsic fluctuations of the network resulted in a certain probability of eliciting the synchronous activation of the neurons (just like in regime 1). The amplitude of such an activation should depend on the level of adaptation the neurons happened to be at, which is at least partly stochastically determined. Indeed, we observed that before the strongest increases in FR (i.e., selecting the highest peaks of the FR around the critical point) the average FR was significantly lower than before low-peak values (Fig. 6A). This state dependence of the FR amplitude suggests that the system is not critical, as in case of, say, self-organized criticality we would expect the amplitude of the avalanches not to depend on the past activity. This phenomenology is not apparent when crossing the boundary separating the bistable and the HAS states (Fig. 6C). Guided by these observations, we contrasted the activity preceding SUT events of low versus high avalanche size in the experimental data. Remarkably, we observed the same state dependence, that is big-size avalanches were preceded by a period of significant lower FR as compared to small-size avalanches (Fig. 6B). The mechanistic underpinnings of this phenomenon results from the key interplay between endogenous fluctuations of neuronal activity *v*(*t*) and the adaptation level *a*(*t*) neurons accumulate (Fig. 6D). If by chance past activity is low and the network is relatively rested, adaptation will be low (Fig. 6D top-right). In a network more excitable like that, firing rate *v*(*t*) can display larger SUTs, as a wider SO cycle can be followed in the phase plane (*a*, *v*) (Fig. 6D, orange traces). The opposite occurs if the network is fatigued (Fig. 6D, green traces). In this case adaptation will be high before a network activation leading to a smaller SO cycle with limited firing rates.

**Figure 6.**
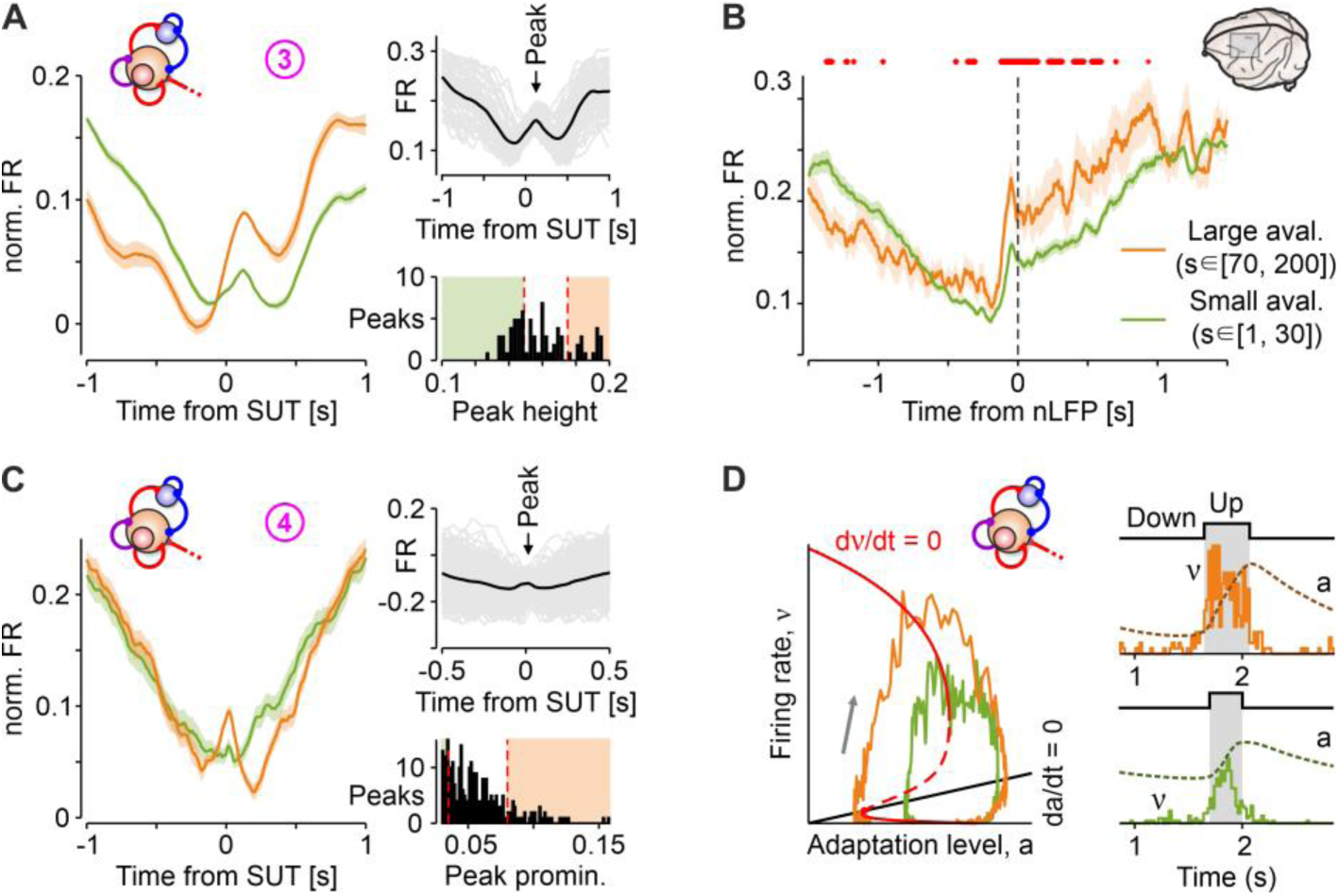
Predicted state-dependency of activations is confirmed in PMd. **A)** Spiking activity at the critical point (regime n.3 as in Fig. 5) for the groups of steepest (orange) and shallowest (green) activations. The waveforms cross each other shortly before the peak, switching from low activity preceding a steeper activation (orange) and from higher activity preceding a shallower activation (green). Insets: normalized FR for all the modules in regime 3 (grey) and their mean (black); Bottom, definition of steepest and shallowest activations used above based on the distribution of the spiking rates at the peak (shown on top panel). **B)** Average normalized FR centered at nLFPs in the PMR epochs corresponding to small (green) and large (orange) *in vivo* avalanche sizes *s*. Red dots: times at which the two average FRs are significantly different (*P* < 0.05). **C)** As in panel A, but for a bistable regime (n.4 as in Fig. 5), in which the network goes back and forth between the asynchronous and the bistable regimes. The inset shows that the peaks are now selected based on their prominence (see Methods). **D)** Sketch of the proposed mechanism explaining the memory effect of population activation. The adaptation level accumulated in the past determines the size of avalanches. When the adaptation level *a* relaxes to lower levels the firing rate can reach higher values (orange), as opposed to the activity jumping to a lower level when *a* is still at intermediate levels (green).

These results show that our simulations reproduced the rich repertoire of dynamical regimes observed in our experiments and predicted the dependence of the amplitude of the slow oscillations from the preceding level of activation of the network. The presence of this memory effect in the cortical data during resting strongly suggests that, while the animals are engaged in the task, the cortical network in the PMR is temporarily transitioning from the awake state to a sleep-like global state.

## Discussion

We have shown for the first time that in resting periods during a behavioral task the premotor cortex undergoes the emergence of spatiotemporal activity patterns assimilable to slow waves as observed in NREM sleep and under anesthesia-induced unconsciousness. This dynamics results from getting close to the critical point separating the desynchronized from the slow oscillatory state, where a synchronization phase transition occurs. As a major novelty with respect to previous works, we characterized this phenomenon relying on the spiking activity of local cell assemblies probed by chronic intra-cortical multi-electrode recordings. The theoretical guidance of a network model of spiking neurons allowed us to predict that the level of activity preceding the slow waves in resting wakefulness shapes their size. An expectation that was confirmed in our experiments, thus strongly supporting the hypothesis that local adaptation of spiking frequency is a key player in approaching the critical state. Hence, the brain of behaving monkeys does not sit in a fixed phase, but rather it explores a landscape of sub-and supercritical states distant from criticality, the latter being occasionally – but repetitively –approached.

The brain has been often associated to a self-organized critical system (SOC), for which criticality is attractive and not regulated by the tuning of an order parameter (Chialvo, 2010; Plenz et al., 2021). Such a system yields scale-free dependence of both temporal and spatial scales and is presumed to offer computational advantages (Bertschinger and Natschläger, 2004; De Arcangelis et al., 2006; Beggs, 2008). Our results contradict this hypothesis in several respects. First, the activity during the task does not show the characteristic signature of criticality (see Figs. 2D and 4E. This is likely due to the fact that spiking activity is highly variable in response to sensory-related inputs posing the network out-of-equilibrium, compatibly with the compelling evidence that motor and premotor cortices visit low-dimensional manifolds (Churchland et al., 2010; Pani et al., 2022). Second, the system does not appear to stay in a fixed phase, but rather it wanders in the space of possible phases, occasionally approaching a critical point separating the desynchronized awake from the synchronized unconscious state. We believe this critical point to be likely repulsive (Pazienti et al., 2022), compatibly with the hypothesis that sleep and wakefulness are two distinct metastable state of the global brain dynamics (Saper et al., 2010; Adamantidis et al., 2019). Instead, our results reinforce with experimental data the hypothesis about the presence of a synchronization phase transition between a desynchronized-conscious state and slow-wave activity associated with unconsciousness (Sanchez-Vives et al., 2017; Di Santo et al., 2018).

One criticism moved to the criticality hypothesis is that power-law distributions of avalanche duration and size – a typical fingerprint of critical scale-free dynamics – could be a sort of side-effect of particularly coarse neural signals – such as the LFP. Our approach does not suffer from these limitations thanks to the use of a measure of the spiking activity targeting local neuronal assemblies. As a matter of fact, we showed instances in which the spiking activity exhibited no signs of criticality in the presence of power-law behavior of the nLFPs (see Fig. 2C and D). This evidence further supports the hypothesis that avalanches of nLFPs often reported in literature (Petermann et al., 2009; Plenz et al., 2021) may confuse truly critical, sub- and super-critical regimes (Touboul and Destexhe, 2010, 2017; Destexhe and Touboul, 2021).

A major result of our work is the observed relationship between the level of activity preceding an avalanche and its size. In the presence of SOC, we would expect any avalanche size to be randomly selected according to the power-law distribution, irrespective of the preceding dynamics of the spiking activity. The “history”-dependence predicted by our network model of spiking neurons arises from the accumulation of fatigue during spike firing. This phenomenon, known since the dawn of neurophysiology as spike-frequency adaptation, dampens the firing rate in a fatigued neuronal system. In this framework, a relatively active network preceding the resting period accumulates more fatigue, thereby increasing the adaptation level. This heightened adaptation ultimately reduces the excitability of the network leading to more localized and brief avalanches (Fig. 6A-B). When the network is relatively inactive during the same period of the task, it becomes less fatigued, leading to the opposite trend eventually facilitating the onset of large slow waves. We believe that such spatiotemporal organization of spontaneous avalanches of activity fundamentally challenges and undermines the SOC hypothesis for cortical networks. This also allows us to rule out another type of critical regime proposed in the literature: the active-to-quiescent transition, which corresponds in our bifurcation diagram to a small region separating awake-like asynchronous activity (HAS) from the bistable phase (Fig. 5A, 6C). This transition is associated with the sand-pile universality class, characterized by random wandering across metastable states (Hahn et al., 2017; Meshulam et al., 2019), and it is known to have significant limitations in accurately describing neuronal recordings (Priesemann et al., 2014).

In addition to this local modulation of adaptation strength over time, our network model incorporates a slow rhythm that influences the excitation level of the examined premotor cortex. This approach aims to replicate the varying levels of network engagement during the execution of reaching tasks, with the pace determined by the sequence of trials conducted throughout an experimental session (Fig. 5). Our theoretical framework is then relatively different from the widely shared picture seeing thalamo-cortical networks as living in nonequilibrium stationary states positioned in a specific location of their bifurcation diagram (Bazhenov et al., 2002; Hill and Tononi, 2005; Destexhe, 2009; Steyn-Ross et al., 2013; Krishnan et al., 2016; Di Santo et al., 2018). Indeed, while these network models exhibit a synchronization phase transition, they do not explore this critical point through alternating perturbations as we have done here. As task execution causes the brain to oscillate near this transition, a hysteresis loop may emerge, as recently observed in an analogous network of chemical relaxation micro-oscillators (Calugaru et al., 2020). This finding would further support previous observations at the mesoscopic scale of local cortical networks, where a subcritical Hopf bifurcation has been characterized in rodents waking from anesthesia (Tort-Colet et al., 2021).

The capability of our network model of spiking neurons to successfully capture the rich phenomenology we observed *in vivo*, confirms that the theoretical compass offered by the excitation-adaptation (E-A) bifurcation plane (Fig. 5) is a powerful tool to predict the dynamical behavior of the premotor cortex across the sleep-wake cycle. In this context, the rhythmic alternation between the unconscious-synchronized and conscious-desynchronized brain states can be represented as a closed trajectory. This representation is similar to previous descriptions of the NREM-REM sleep cycle (Steyn-Ross et al., 2005). It also parallels the cyclic alternating pattern of slow oscillations and micro-arousals (Terzano et al., 1985; Parrino et al., 2012), signaling an unstable phase of the unconscious state (Tort-Colet et al., 2021). The force driving the brain along the sleep-wake trajectory must be directly related to the process S postulated by (Borbély, 1982) and lately observed both in humans (Achermann and Borbély, 1997) and rodents (Thomas et al., 2020). The process S is associated to sleep pressure accumulating during wakefulness, including in our case the time when the monkeys are engaged in performing the task, and lowering during sleep (Fig. 7A). In the E-A plane, process S can be modeled as a drifting force moving the system towards the SO phase (strong adaptation) and back to the awake-like phase (HAS, relatively weak adaptation). Sleep pressure brings the brain closer to the critical point that separates awake and sleep-like states (see Fig. 7B-left, brown section of the cycle). If this was the case, at the beginning and at the end of an experimental session we would expect the adaptation strength in the premotor cortex to be different. Specifically, we would expect larger and more pronounced waves to spontaneously emerge in cortical activity during rest and, at the same time, deeper (larger) nLFPs. This in turn would result in spindle oscillations with greater power near the slow oscillation (SO) phase, rather than at the beginning of the session when sleep pressure is weaker. Remarkably, both these expectations have been confirmed in our experiments (see Fig. 7B and Fig. 3C). Thus, sleep pressure and local fatigue can be seen as components of a restless system wandering in the E-A plane, close to a critical point of brain dynamics. This may also impact behavioral performance, potentially leading to lapses in attention due to more frequent intrusions of slow waves—essentially a form of local sleep activity—in the awake brain. This phenomenon has recently been observed in humans as nLFPs from scalp EEG (Andrillon et al., 2021). If confirmed, this evidence would challenge the conventional belief that criticality enhances cognitive processing.

**Figure 7.**
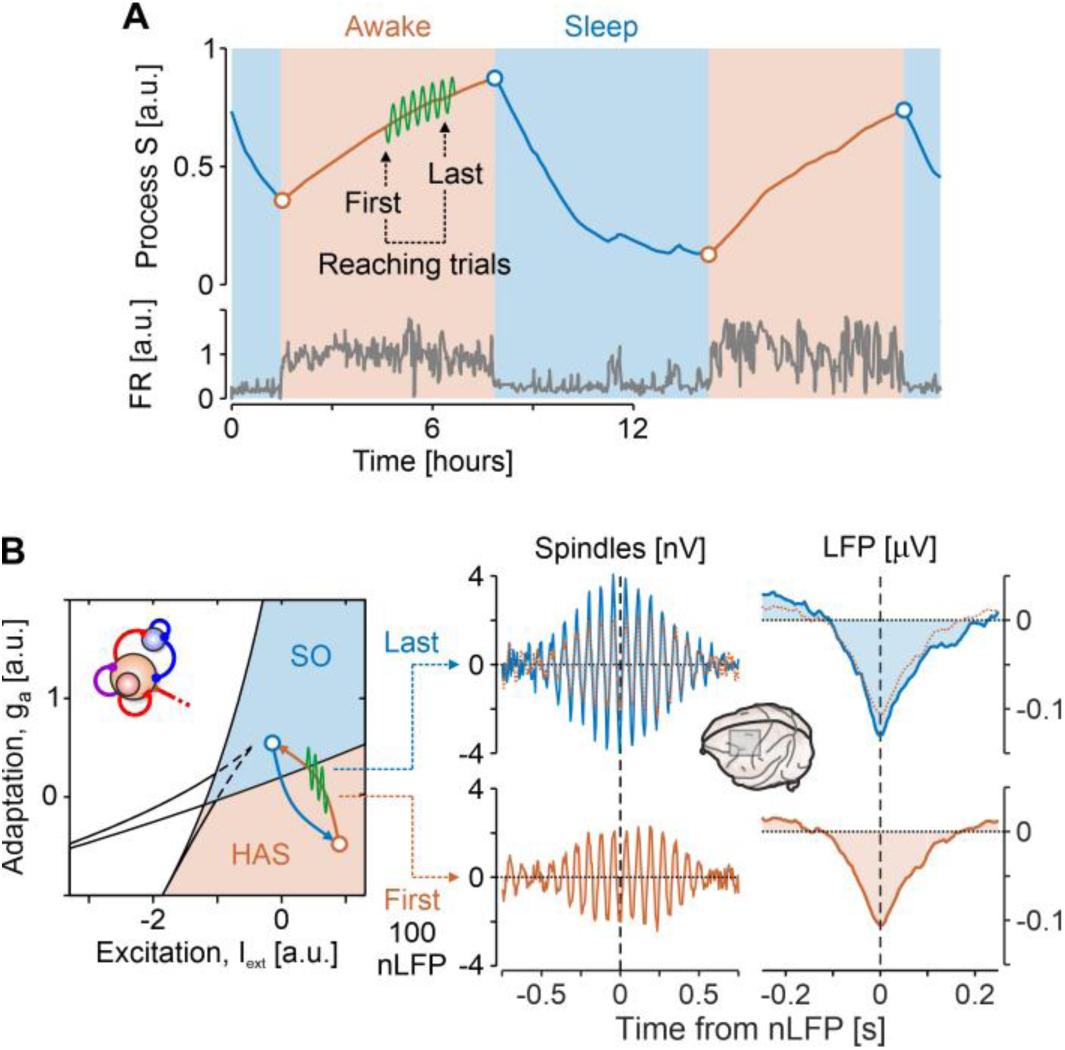
Sleep pressure pushes the PMd to the edge of synchronization. **A)** According to the theories of sleep homeostasis a process S (top) determines the pressure to sleep (Borbély, 1982): accumulates during wakefulness (orange) and reduces during sleep (blue). Circles, sleep-wake transitions. Green oscillations, modulation of excitation in network models expected while performing a behavioral task (see Fig. 5). Bottom, firing rate from a neuronal assembly [adapted from (Thomas et al., 2020)]. **B)** In the (excitation, adaptation) plane (left) the process S drives the cortical network to oscillate back and forth between the Sleep- and Awake-like phases (blue and orange regions as in Fig. 5A). First and last 100 nLFP singled out in Monkey P during PMR represent the beginning and the end the reaching task session (top and bottom panels on the right). Spindles are the band-pass filtered LFPs centered around nLFP and averaged in the first and last 100-group (blue and orange, respectively). Right, nLFP averaged across the same groups.

## Acknowledgements

Work partially funded by EU H2020 Research and Innovation Programme, Grant 945539 (HBP SGA3) and by the Italian National Recovery and Resilience Plan (PNRR), M4C2, NextGenerationEU (Project IR0000011, CUP B51E22000150006, ‘EBRAINS-Italy’) to MM and SF.

## Materials and Methods

### Experimental setting

#### Subjects

Two male rhesus macaque monkeys (Macaca mulatta; designated Monkeys P and C), weighing 6 and 9 kg, were studied. The monkeys were pair-housed with cage enrichment. They were fed daily with standard primate chow, supplemented with nuts and fresh fruits. The monkeys received part of their daily water supply during the experiments in the form of fruit juice. All experimental procedures, animal care, housing, and surgical procedures conformed to European (Directive 2010/63/UE) and Italian (D.L. 26/2014) laws on the use of nonhuman primates in scientific research and were approved by the Italian Ministry of Health.

#### Animal preparation

At the end of training period a Utah array (96 channels, Blackrock Microsystems, USA) was surgically inserted in the PMd of each monkey, using the arcuate sulcus (AS) and pre-central dimple (pCD) as anatomical landmarks after opening of the dura (Fig. 1 A for Monkey C). The recording site was contralateral to the arm used during the experiment. A head-holding device was implanted in both monkeys. All surgeries were performed under sterile conditions and veterinary supervision. Antibiotics and analgesics were administered postoperatively. Anaesthesia was induced with ketamine (Imalgene, 5-10 mg kg−1 i.m.) and medetomidine hydrochloride (Domitor, 0.03-0.04 mg kg−1 i.m.) and maintained with inhalant isoflurane (0.5%–4%) in oxygen. Antibiotics were administered prophylactically during the surgery and postoperatively for at least 1 week. Postoperative analgesics were given at least twice daily. Recordings of neuronal activity started after a minimum of 10 weeks after recovery from the surgery.

#### Apparatus and task

Experiments were performed in a darkened, acoustically insulated room. Monkeys were seated in front of a black isoluminant background (<.1 cd/m2) of a 17-inch touchscreen monitor (LCD, 800 × 600 resolution). A non-commercial software package, CORTEX (www.nimh.nih.gov), was used to control the presentation of stimuli and behavioural responses.

The two monkeys performed a standard delayed reaching task (Fig. 1A right; (Weinrich and Wise, 1982; Crammond and Kalaska, 2000)), where Go signal occurred after a random delay (800–1600 ms) since the target appearance (either left or right side of the screen on different trials).

At the end of correct trials, the monkeys experienced a brief sound that was accompanied by the delivery of juice as a reward. In signal-respond trials (errors), neither sound nor reward was delivered, and the screen turned blank. The intertrial interval was set to 1000 msec.

Anesthesia-induced slow oscillations were obtained in both animals by ketamine (Imalgene) 25 mg/kg i.m., in separate sessions of those used for task recordings.

#### Neural recordings

Unfiltered raw activity was recorded from 96 channels using a Utah array (Blackrock Microsystems, USA) and a TDT System 3 (Tucker Davies Technologies, sampling rate 24.4 kHz).

### MUA estimate and state transition detection

Up and Down states and spectral-derived multi-unit activity (referred to as spiking activity or normalized firing rate, FR) were estimated from the recorded raw signals (Mattia et al., 2010; Reig et al., 2010; Sanchez-Vives et al., 2010). Briefly, the power spectra were computed from 5-ms sliding windows of the raw signal. spiking activity was estimated as the relative change of the power in the [0.2, 1.5] kHz frequency band. Such spectral estimate of the spiking activity was not affected by the electrode filtering properties, and it provided a good estimate of the relative firing rate of the pool of neurons near the electrode tip (Supèr and Roelfsema, 2005; Stark and Abeles, 2007). In a second step, the spiking activity was logarithmically scaled to compensate for the high positive fluctuations due to the neurons closest to the electrode, thus obtaining the spiking activity signal. We then smoothed the spiking activity by performing a moving average with a sliding window of 40 ms. Reference value for this signal was associated to the most prominent peak of its distribution corresponding to the average activity during Down states and set to 0. From the long-tailed histogram of the spiking activity, an optimal threshold separating Up and Down activity states was set at the absolute value of 0.4. The threshold for detecting both Down-to-Up transitions was the same. For each dataset, we checked the quality of the channels and excluded from further analysis the ones where the signal-to-noise ratio was particularly weak. The number of channels artifact free that were analyzed was 96 (43) for the delayed-reaching experiment and 59 (64) for the experiment in anesthesia for monkey P (C).

The frequency of the SO was the inverse of the duration of the entire Up-Down cycle. We reconstructed the activation waves from the detected Down-to-Up state transitions as in (Capone et al., 2019; Dasilva et al., 2021; Pazienti et al., 2022). Briefly, each wave was reconstructed pooling together the transitions occurring in multiple electrodes in a reasonable time interval that was iteratively reduced until each wave contained no more than one transition per channel. Each reconstructed wave was associated with a vector of relative time lags computed as the difference between the time of occurrence of the wave in each electrode and the average time of occurrence across all the electrodes taking part in the wave propagation. We rejected the waves occurring in less than twelve channels, otherwise we replaced the missing values with the result of a nearest-neighbor interpolation using the five nearest points in terms of Euclidian distance of wavefronts/time lag vectors. Using the resulting vectors describing the detected activation wavefronts, for each animal and anesthesia level we composed a time-lag matrix (TLM) having the number of rows equal to the number of waves and the number of columns equal to the number of recording channels of the experiment. We refer to the rows of the TLM, representing relative Down-Up transitions for each channel, also as time-lag arrays. The spatiotemporal course of the wavefronts was then obtained by spatially interpolating the time lags without smoothing, using a thin-plate spline method.

### nLFP extraction

Local field potentials (LFP) were calculated as the components of the recorded signals in the frequency band [1, 200] Hz. High- and low-pass filtering of the original (raw) signal was achieved by first down sampling into 5 ms time bins and then subtracting from the outcome the component below 1 Hz. Finally, a smoothing kernel was applied with a characteristic time scale of 40 ms.

Negative local field deflections (nLFP) were computed as the time points in which the LFP exceeded a negative threshold. The threshold was automatically calculated as the standard deviation of the LFP multiplied by a factor; the latter was chosen to be equal to 2.0.

#### Relevant intervals of interest

The nLFP-centered analysis was performed on instances in which both the following conditions were fulfilled: the nLFP distribution showed a maximum and the spiking activity a minimum (i.e., it was close to zero). As it turned out that these conditions were fulfilled after the second movement completion in the task (Mvt2 in Fig.1 D) these time intervals are referred to in the text as Post Movement Rest (PMR).

For monkey P, this corresponded to the region just after the end of the Mvt2. Specifically, we stored all nLFPs occurring in the interval [0.5, 2] s after the related marker set by the experimenters. For monkey C, the 1.5 s-interval selected was the inter-trial period, i.e. all the time segments between the end of the trial and the beginning of the next one, which occurred slightly later than Mvt2.

#### Channels participation

In order to evaluate the participation of single channels to the nLFP statistics (as to produce Fig. 1B, bottom), the same procedure was applied as described above, however on a single-channel basis.

### Avalanches computation

Neuronal avalanches were computed following the procedure illustrated in (Petermann et al., 2009). Briefly, signal events (either nLFP or SUT occurrence) were binned and each avalanche event was defined as the cumulative number of occurrences until an empty bin was reached. To rule out a systematic bias owing to the choice of time bin, we employed the same heuristic prescription as that of (Beggs and Plenz, 2003), namely to create a pooled time series with activations (nLFP or SUT) from all electrodes, and to use as time bin the average inter-event interval (IEI, see Fig. 1b of (Ribeiro et al., 2010)), that is the time between consecutive activations (whether or not from the same electrode).

Figs. 2C and 2D contrast avalanche statistics during low spiking/high nLFP density as defined above (after the reaching movement for monkey P and in the inter-trial intervals for monkey C) versus the distribution of nLFPs and FR of the whole trial.

In order to estimate the FR preceding avalanches of different size (as for Fig. 6B) the average FR was computed by restricting the avalanche computation to two distinct and non-overlapping sizes *s*: in the intervals *s* ∈ [1,30] and *s* ∈ [70,200] for monkey P and *s* ∈ [1,15] and *s* ∈ [35,200] for monkey C.

### Estimate of SUT times

For the experimental data recorded during the task, in order to compute the matrices of the SUT times per channel and per nLFP event, as the signal-to-noise ratio was relatively small, we opted to fit the waveforms around the SUT to get estimates that were more robust. To do so we followed three steps: i) we computed the cross-correlation between the average nLFP-centered spiking activity transition and all individual FR ones in the time window 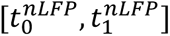 and selected only the ones significantly correlated; ii) we then computed, for each trigger event (i.e., nLFP event), the cross-correlation between the FR transition in the same time interval 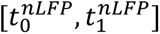 and the corresponding transition across all channels, and selected only the significant transitions at the channels level; iii) for each channel and each transition, we performed a sigmoidal fit to estimate the exact time the waveform was undergoing the SUT. 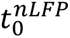 and 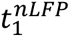 were chosen to be [−0.09 *s*, +0.06 *s*] for monkey C and [−0.20 *s*, −0.04 *s*] for monkey P.

The sigmoidal fit aimed to fit the SUT transitions with the following formula:

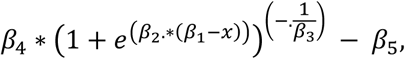

i.e., to find the values *β*_*i*_, *i* = 1,2, …, 5 for each SUT transition. For this purpose, the MATLAB function `nlinfit` was used with initial values equal to ***β*** = [−0.05, 100, 1, 0.5, 0.3]. Once the fitting procedure produced an outcome, the minimum σ_*min*_ and maximum σ_*max*_ values of the resulting fit were extracted, and the time of transition was set as the time when the resulting sigmoidal function trespassed the threshold 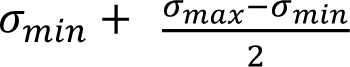 (i.e., half the height of the sigmoidal function). In case no threshold crossing was detected, the fitting procedure was re-applied on the waveform shifted to the left or to the right by *x* bins, where *x* ∈ [−*A*, *A*] and *A* was either 10 or 15 bins. After that, time of threshold crossing was estimated for each shift, and the shift that yielded the minimum difference between the original waveform and the fit was selected. If none of the previous iterations yielded a time of threshold crossing, the SUT time for this channel and this nLFP was not assigned any value and did not contribute to the time matrix.

### Traveling wave detection and reconstruction

Relying on the first principal component of TLM we clustered waves/rows sharing similar features with a k-means algorithm. The optimal number of clusters to be used to group the waves in each anesthesia level was chosen using a Silhouette method. For each cluster, we obtained the mean wavefront and its spatiotemporal profile by averaging the waves belonging to it. The mean wavefront extracted from each cluster was used to compute the mean velocity and direction of propagation of the cluster as in (Capone et al., 2019; Pazienti et al., 2022). This was done relying on the smoothed surface *T*(*x*, *y*) (thin-plate smoothing spline) of the relative time lags associated with each cluster and distributed according to the electrode positions, and computing the local velocity as *V*(*x*, *y*) = 1/(∂_*x*_*T*^2^ + ∂_*y*_*T*^2^). The gradient of this field points out the direction of the wave propagation. Mean values are computed across the used electrode positions.

### Snapshots of activity on the grid

In order to show the average Down-to-Up transition every channel experienced and the corresponding wavefront on the recording grid of electrodes (Figs. 3B and 4C-D), we proceeded with the following steps. We normalized every average waveform by subtracting the baseline (defined as the activity preceding 175 ms before the corresponding nLFP) and dividing it by its standard deviation (SD).

The corresponding time of propagation per channel were extracted as the relative time trespassing a threshold set at 1 SD. Contours of propagation were computed by thin-plate smoothing splines using the Matlab function `tpaps` with smoothing parameter equaling 0.8. Finally, the filled areas shown in panels 3B, and 4C,D were obtained by simply filling all the array surface where activity had already crossed the threshold at the corresponding time.

For the snapshots of the simulated activity on the grid shown in Fig. 5C (bottom) we applied a spatial cubic smoothing spline (computed with the Matlab function `csaps` with smoothing parameter set to 0.8) to the simulated FR and superimposed it with the contour of the activity at a given level (white thick line).

### Models and simulations

Network model shown in Figure 5 was composed of standard point-like integrated-and-fire (LIF) neurons. It is organized as a set of interacting modules, each consisting of two interconnected pools of excitatory and inhibitory spiking neurons. Similarly to (Mattia et al., 2013), each cortical module was composed of 250 inhibitory (I, 20%) and 1,000 excitatory (E, 80%) neurons with spike frequency adaptation (SFA). The neurons in the excitatory population were divided into a “foreground” (F, 25%) and “background” (B, 75%) sub-pools. Such modules were spatially distributed on a 2-dimensional grid of 13 × 13 sites. The inter-modular connectivity was greater than zero only between pairs of populations displaced at relative distance less than 3, thus modelling cortico-cortical horizontal connections. Membrane potential *V*(*t*) of each neuron evolved according to *V̇*(*t*) = −*V*(*t*)/*τ* + *I*(*t*) − *a*(*t*) where 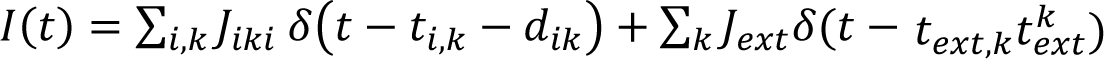 is the synaptic current from recurrent (*i*) and external (*ext*) presynaptic neurons emitting their *k*-th at time *t_i,k_* and *t_ext,k_*, and transmitted with efficacy *J_i_* and *J_ext_*, respectively. Recurrent spikes are delivered with axonal transmission delay *d_i_*. The adaptation current 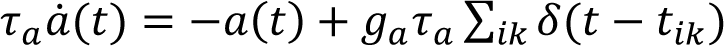 models the activity-dependent afterhyperpolarizing *K*^+^ current underlying SFA, which increases by *g_a_* = 0.04 mV/ms at each spike emission time *t_k_* of the neuron and relaxing with a characteristic time *τ*_*a*_ = 500 ms. No SFA was incorporated in inhibitory neurons. Membrane potential decay constants were *τ* = 20 and 10 ms for E and I neurons, respectively. Spikes were emitted when *V*(*t*) crossed the threshold Ɵ = 20 mV, after which potential was reset to 16 mV for an absolute refractory period *τ*_0_ = 2 and 1 ms for E and I neurons, respectively. External spikes mimicked the input from the other cerebral areas and were implemented as a Poisson process from *C_ext_* independent source firing at rate *v_ext_*. We choose *C*_*ext*_ = 800, *v*_*ext*_ = 3 *Hz*, *J*_*ext*_ = 0.43 mV and 0.56 mV for E and I neurons, respectively, in order to have a fixed point with firing rate for E and I neurons at *v_E_* = 3 Hz and *v*_*I*_ = 6 Hz, respectively, under mean-field approximation (Amit and Brunel, 1997). To this purpose the synaptic efficacies were randomly chosen from a Gaussian distribution with mean *J_αβ_* and standard deviation Δ*J_αβ_* depending on the type of presynaptic (*β* = *E*, *I*) and postsynaptic (α = *E*, *I*) neurons, whereas Δ*J_αβ_* = 0.25 *J_αβ_* for any *α* and *β*. Inhibitory connections were set to *J*_*aa*_ = *J*_*Ea*_ = −1.5 mV, *J*_*EE*_ = 0.43 mV and *J*_*IE*_ = 0.56 mV. In the same mean-field framework we structured the cortical module to have an additional high-frequency fixed point for the F excitatory neurons by potentiating synaptic self-excitation *J_FF_* and depressing the other excitatory connections *J_FBV_* and *J_BF_* compared to *J_EE_* as in (Amit and Brunel, 1997): *J_FF_* = 0.61 mV and *J_FB_* = *J_BF_* = 0.39 mV. Intra-modular connectivity between excitatory neurons (i.e., the probability be synaptically coupled) was *C_EE_* = 0.8, whilst the inter-modular connectivity decreased with distance (Ercsey-Ravasz et al., 2013) assuming the values (0.03, 0.018, 0.007, 0.004, 0.001, 0.001). Axonal transmission delays were drawn from an exponential distribution with mean 〈*d*_α*E*_〉 = 22.6 ms and 〈*d*_α*a*_〉 = 5.7 ms (α = *E*, *I*). In the 13 × 13 grid, the modules in the two opposite 2 × 2 corners shown in Fig. 5A have an enhanced self-excitation among F neurons (*J_FF_* = 0.65 mV) to increase their excitability. Analyses were performed in the 5 × 5 subset of modules in the center of the grid. To use the same analytical approach developed for *in vivo* recordings, we transduced the simulated activity in an *in silico* representation of the spiking activity. The latter was computed from the firing rates *v_F_*(*t*) of each module, subsampled at 5 *ms*, by adding a white noise *ζ*(*t*) with 〈*ζ*(*t*)〉 = 0 and 〈*ζ*^2^(*t*)〉 = *η*^2^, where *η* = 0.4 · min (*v_F_*(*t*)), to mimic the fluctuations observed in the experiments.

Simulations were performed with NEST (Gewaltig and Diesmann, 2007). Full Python code where all the missing model parameters and information is freely available in EBRAINS Knowledge Graph (https://search.kg.ebrains.eu/instances/Model/eb925263fd703c9c7aec2ee94e8be72520e3b69e).

